# Cultivation of plant-growth promoters in vinasse: contributions for a circular and green economy

**DOI:** 10.1101/2022.12.28.522132

**Authors:** Mariela Analía Torres, Alejandra Leonor Valdez, María Virginia Angelicola, Enzo Emanuel Raimondo, Hipólito Fernando Pajot, Carlos Gabriel Nieto-Peñalver

## Abstract

Vinasse is a by-product with a key role in the circular economy. In this work, we analyze sugarcane vinasse as culture medium for obtaining single and mixed inoculants. *Trichoderma harzianum* was cultured in single and sequential co-culture with *Pseudomonas capeferrum* or *Rhizobium* sp. Fungal biomass was higher in vinasse than in a laboratory medium. Residual vinasses presented almost neutral pH and lower conductivities and toxicity than raw vinasse. Fertigation with residual vinasses improves characteristics of soil evidenced in the total N, cation exchange capacity, urease and acid phosphatase, and the microbial metabolic diversity, in comparison to raw vinasse. The evaluation of the treatment indicates that vinasse is suitable for the production of inoculants containing *T. harzianum* and that the co-culture with *P. capeferrum* improves the characteristics of the residual vinasse in comparison to *Rhizobium* sp. Obtaining this valuable biomass in vinasse is relevant for the circular and green economy.

## 1. Introduction

The main effluent generated after ethanol production by the sugar-alcohol industry is vinasse, a dark brown liquid with an acidic pH (3.5-5.0), and high Biological and Chemical Oxygen Demands (54,000 mg L^−1^ and 110,000 mg L^−1^, respectively). Nearly 13 litres of vinasse are generated on average for each litre of ethanol produced, meaning 13·10^6^ L per year for an average refinery (Christofoletti et al., 2013). Vinasses consist of 93% water and 7% solids, with a high prevalence of potassium, phenolics and melanoidins (de Godoi et al., 2019). Together with the acid pH and the volume, these are the main responsible for the negative ecological impact when vinasses are not properly disposed (Christofoletti et al., 2013). Different strategies have been envisaged in order to diminish its environmental hazard. Fertigation, i.e., its disposal in soils as fertilizer, is one of the most utilized methods for its management. This practice allows the recycling of water, organic matter and important elements, including nitrogen and phosphorus. However, excessive or improper fertigation leads to the acidification, salinization, and the reduction of microbial activity in affected soils (Christofoletti et al., 2013).

Several reports analyzed the microbial treatment of vinasse. In these cases, the microbial metabolism consumes the organic compounds and converts them to CO_2_ and water, reducing the toxicity of vinasse (Chuppa-Tostain et al., 2020; España-Gamboa et al., 2011). The main drawback for the effective application of these bioremediation processes are the enormous volumes of vinasse generated in a daily basis. Other approaches analyzed the utilization of vinasse as culture medium for producing microbial biomass (Candido et al., 2022; Montalvo et al., 2019), metabolites (Altenhofen da Silva et al., 2017) or enzymes (Ahmed et al., 2022). Interestingly, the production of particulate fertilizers in vinasse has been recently reported (Cerri et al., 2020). This strategy clearly shows that vinasse is not necessarily a waste but a by-product. In addition, when vinasse is utilized instead of other sources for the production of these metabolites or for obtaining these biomasses, it becomes a highly valuable by-product with a key role in the future of circular economy (Hoarau et al., 2018; Karp et al., 2022). Circular economy proposes a cyclical production system in which ‘wastes’ are considered supplies to next processes in a cyclical productive system (Kneese, 1988). The actual and current relevance of circular economy is evidenced in the manner the Sustainable Development Goals of the United Nations are traversed by this concept. At the same time, the concept of ‘green economy’, i.e., the development of biotechnological processes of economical relevance in a sustainable manner, has also to be taken into account.

In this work, we evaluated vinasse from sugarcane as culture medium for the production of the plant-growth promoter *Trichoderma (T.) harzianum* MT2, in single and in co-culture with *Pseudomonas capeferrum* WCS358 and *Rhizobium* sp. N21.2. *T. harzianum* MT2 is a native isolate obtained from tomato rhizosphere (Malinar, 2020). *T*. *harzianum* is a relevant biological control agent that promotes plant growth through the mycoparasitism of fungal phytopathogens. In addition, *T. harzianum* also stimulates root and shoot elongation, the uptake of nutrients and increase the stress resistance. *T*. *harzianum* and related species are at the top of the formulated fungal biofertillizer in a global market estimated at U$S1.66 billion by 2022 (Aloo et al., 2021). Mass production of *T. harzianum* is obtained in solid-state fermentations employing seeds of rice, sorghum, and agro-waste products as substrate, or in liquid fermentations in a wide variety of culture media (Dutta et al., 2022).

Mixed inoculants show synergistic potentials for promoting plant growth in contrast to single inoculants (Santos et al., 2019). Mixed bioinoculants containing *Trichoderma* spp. usually include *Bacillus* spp., *Pseudomonas* spp. (Poveda and Eugui, 2022), and rhizobia (Barbosa et al., 2022), among other bacteria. The simplest way of producing mixed inoculants consist in the cultivation of individual strains in axenic cultures that will be mixed in the final commercial product. For a large scale development, this represents an extra cost of production. Vinasse offers the possibility for obtaining mixed inoculants, as reported previously for other biotechnological applications (Eder et al., 2020; Iltchenco et al., 2020). The simultaneous production of mixed inoculants in vinasse as substrate may present the advantage of generating less-toxic wastes. In co-culture with *T*. *harzianum* MT2, we analyze in this article two model bacteria with plant-growth promoting potential. *P. capeferrum* WCS358, isolated from potato rhizosphere (Geels and Schippers, 1983), induces the systemic resistance of host plants and secretes pyoverdine siderophore that inhibits phytopathogens (Berendsen et al., 2015). *Rhizobium* sp. N21.2 is a native isolate previously isolated from strawberry rhizosphere. N21.2 strain produces siderophores, solubilizes phosphates, secretes auxins and putatively fixes nitrogen, according to positive PCR amplification of the nitrogenase *nifH* gene (unpublished results).

The aim of this work was to analyze the utilization of the vinasse as culture medium for the production of *T*. *harzianum* MT2, in single and in co-culture with two model plant-growth promoting bacteria. As a contribution to the circular and green economy, the objective is to obtain valuable biomass in a cost-effective manner from an abundant by-product, reducing at the same time the environmental impact of the residual vinasses.

## 2. Materials and Methods

### 2.1 Strains and culture media

*Trichoderma* (*T*.) *harzianum* MT2 was isolated from tomato rhizosphere and identified after sequencing the ITS region and the D1/D2 region. *T*. *harzianum* MT2 was maintained in Yeast-Malt Extract medium (YME) medium (Tavares et al., 2005). *Pseudomonas* (*P*.) *capeferrum* WCS358 was maintained in Luria Bertani (LB) medium. *Rhizobium* sp. N21.2 was maintained in Yeast Mannitol (YMA) medium (Vincent, 1970). All strains were preserved at −80 °C in 20% glycerol.

### 2.2 Source and characterization of vinasse

Vinasse obtained directly from the distillery columns was provided by a local sugarcane biorefinery located in Cruz Alta, Tucumán, Argentina during the sugarcane harvest in 2019, and stored at −20 °C. Vinasse was physically and chemically characterized at Estación Experimental Agroindustrial Obispo Colombres (EEAOC, Tucumán, Argentina), following standard procedures: fixed and volatile solids by the Standard Method 2540E (Fixed and Volatile Solids Ignited at 550°C); total solids by the Standard Method 2540C (Total Dissolved Solids Dried at 180 °C); pH and conductivity by the Standard Method 4500 and 2510 with a pH meter and conductivity meter, respectively; chloride by the Standard Method 4500-Cl--B and sulfide by the Standard Method 4500-S_2_^−^-D; Total Phosphorus by the Standard Method 4500-P-C; Brix by the refractometric method; organic material by gravimetry; Potassium, Calcium, Magnesium and Sodium by Flame Atomic Emission Spectroscopy; total Nitrogen by Kjeldahl method (Suppl. Table 1). Before utilization, the vinasse was centrifuged at 10.000 *g* for 5 min to discard the coarse sludge and then autoclaved.

### 2.3 Fungal and bacterial tolerance to vinasse

Before analyzing the respective growth, the tolerance to vinasse (i.e., the maximal concentration of vinasse that supported the microbial growth) of each strain was determined. Overnight precultures were utilized to inoculate flasks containing 10, 20, 30, 40, 50, 75 and 100% vinasse (dilutions in distilled water). *T. harzianum* MT2 was incubated at 25 °C and 250 rpm. At 0, 48 and 72 h of incubation, dry weights were measured after drying the cell pellets at 105 °C. *P. capeferrum* WCS358 and *Rhizobium* sp. N21.2 were incubated at 30 °C in an orbital shaker at 180 rpm. After 0, 48 and 72 h of incubation, values of CFU mL^−1^ were determined after plating serial dilutions in LB agar or YMA agar for *P. capeferrum* WCS358 and *Rhizobium* sp. N21.2, respectively. For comparison, tolerance values were expressed as percentages considering 100% the values obtained after the addition of the corresponding inocula.

### 2.4 Fungal and bacterial single cultures in vinasse

Microbial growths were evaluated at subinhibitory concentrations of vinasse determined as described above. *P. capeferrum* WCS358 and *Rhizobium* sp. N21.2 were grown in diluted vinasse (10 and 20%) and compared with growth in LB and YMA broth, respectively. Cultures were performed aerobically at 30 °C and 180 rpm. Samples were withdrawn during incubations, and the bacterial growths were evaluated through the determination of CFU mL^−1^ after plating serial dilutions in LB agar and YMA agar. *T. harzianum* MT2 growth was evaluated in 10 and 50 % vinasse at 25 °C and 250 rpm and compared to that in YME broth. Samples were also periodically withdrawn, centrifuged and the cell pellets were dried at 105 °C for the evaluation of the growth by dry weight measurement.

### 2.5 Sequential co-cultures of *T. harzianum* with bacteria in vinasse

Sequential co-cultures of *P*. *capeferrum* WCS358+*T. harzianum* MT2 and *Rhizobium* sp. N21.2 +*T. harzianum* MT2 were carried out as follows. First, *P*. *capeferrum* WCS358 and *Rhizobium* sp. N21.2 were independently cultured in 10 % vinasse for 48 h at 30 °C and 180 rpm. *T. harzianum* MT2 was then inoculated from a 48 h preculture, and pure vinasse was added to achieve a final concentration of 50% (Suppl. Fig. 1). Incubation was continued at 25 °C and 250 rpm for another 72 h. Bacterial and fungal growths were evaluated through the determination of CFU mL^−1^ and the dry weight method, respectively. Single bacterial and fungal cultures were carried out as controls. At the end of the incubations, complete biomasses were removed by centrifugation at 10.000 *g* for 10 min and the residual vinasses were analyzed as described below (Suppl. Fig. 1).

### 2.6 Physical-chemical analysis and toxicity of vinasses

Acidity was measured with a pHmetrer (Sartorius). Dissolved solids were measured with a brixometer (Arcano) and conductivity was determined with a conductimeter (COM-100, HM Digital). Toxicity of residual vinasse was evaluated by the acute toxicity assay with *Lactuca sativa* seeds. Briefly, seeds were placed in Petri dishes, and 5 mL of residual vinasses or control vinasse (i.e., 50% pure vinasse) all previously diluted in water (1:5) were added. Seeds were also treated with water as control. Plates were incubated in darkness at 25 °C for 5 days. The percentages of germination were determined considering 100% the number of seeds germinated with water. The hypocotyl and radicle lengths were measured and values were compared to those obtained with control vinasse.

### 2.7 Soil fertigation with residual vinasse

Soil not previously fertigated was collected from a local sugarcane farm. Samples were taken from the first 10 cm depth in the furrow zone and nearby the plants. Soil was sieved and then placed in trays for fertigation, as follows. Residual vinasses from single and co-cultures were filtered with gauze to discard the coarse fungal biomass and utilized for the fertigation by aspersion on days 0 and 7 applying 1 L m^−2^ (irrigation sheets=1 mm) in agreement with local recommendations for fertigation with vinasse (Morandini and Quaia, 2013). A group of trays was irrigated with control vinasse (i.e., 50% pure vinasse) or water as control treatments. All trays were incubated at 25 °C for a total of 14 days, and then soil samples were taken for physical-chemical and biological characterizations. Water contents were daily adjusted to the initial values with distilled water considering the loss of weight.

### 2.8 Physical-chemical and biological characterization of soil samples

Soil pH and conductivity were determined in at least three independent samples by mixing one volume of soil was mixed with one volume of distilled water and mixed vigorously. After decantation, pH and conductivity were measured in the soil slurries with a pHmeter (Sartorius) and a conductimeter (COM-100, HM Digital), respectively. Soil toxicity was determined in the same soil slurries with lettuce seeds as described previously. For other chemical characteristics, samples of soils from three independent assays were pooled before analysis: Carbonate content by calcimetry; Total Organic Carbon (TOC) by the Walkley-Black method; total Nitrogen by the Kjeldahl method; available Phosphorus by the Bray-Kurtz method; Cation Exchange Capacity (CEC) was calculated from the quatification of exchange cations (Ca^2+^, Mg^2+^ and K^+^), which were determined by the ammonium acetate method, except for Na^+^ that was determined by the Saturated Paste method. Soil enzymatic activities were determined in at least three independent samples as follows. Hydrolysis of fluorescein diacetate (FDA; μg fluorescein g^−1^ h^−1^), acid phosphatase (AP; μg *p*-nitrophenol g^−1^ h^−1^) and urease (UA; μg N-NH_4_ g^−1^ h^−1^) activities were quantified by spectrophotometry employing published protocols (Adam and Duncan, 2001; Jastrzębska, 2011), with modifications (Raimondo et al., 2019). Catalase activities (CAT; mmol H_2_O_2_ consumed g^−1^ h^−1^) were determined by titration (Jastrzębska, 2011), with modifications (Raimondo et al., 2019). Quantification of heterotrophic microorganisms was performed in at least three independent samples in Plate Count Agar (PCA) by plating serial dilutions of soil supernatants on PCA in triplicate and incubating at 30 °C for 120 h. Results were expressed as CFU g^−1^ of soil. Microbial functional diversity was analyzed by studying patterns of carbon source utilization with Biolog EcoPlates. Microorganisms were extracted from 5 g of soil from each treatment after shaking at 200 rpm at 25 °C for 45 min with 45 mL of 0.9 % NaCl solution. The coarse particles were allowed to decant for 30 min at room temperature and 150 μL of these suspensions were utilized to seed wells of Biolog EcoPlates. Microplates were incubated at 25 °C for 7 days and the absorbance was measured on a daily basis at 590 nm (A_590_). The average metabolic response (AMR) for each treatment was calculated as the mean difference between the A_590_ of wells containing a carbon source (A_590CS_) and the control well with water (A_590W_): AMR=Σ(A_590CS_-A_590W_)/95 (Konopka et al., 1998). Principal Component Analysis was performed with data obtained after 72 h of incubation, when larger differences in the AMR among treatments were determined.

### 2.9 Statistics analysis

All assays were performed independently at least in triplicates. Vinasses were analyzed with ANOVA followed by Dunnett’s post hoc test at *P*<0.05 considering the control vinasse as control.

Conductivity, pH, toxicity, counts of heterotrophic microorganisms and enzyme activities in soil were analyzed with ANOVA followed by Dunnett’s post hoc test at *P*<0.05 considering the soil treated with control vinasse as control. Statistics were performed with Minitab 19 software.

## 3 Results

### 3.1 Vinasse as growth medium for single cultures

Growths of *P*. *capeferrum* WCS358, *Rhizobium* sp. N21.2 and *T. harzianum* MT2 were evaluated in subinhibitory dilutions of vinasse. Tolerance determinations showed that 30% vinasse was inhibitory for both bacteria (Suppl. Fig. 2a and b). *Rhizobium* sp. N21.2 and *P*. *capeferrum* WCS358 growths were then evaluated in 10% and 20% vinasse, and compared to YMA and LB broth, respectively. *Rhizobium* sp. N21.2 showed an 8 h-lag phase in 10% vinasse and then attained 3.03·10^8^ CFU mL^−1^, lower than in YMA medium (1.90·10^9^ CFU mL^−1^) (Fig. 1a). In contrast, CFU mL^−1^ decreased after 8 h of incubation in 20% vinasse, reaching 7.03·10^3^ CFU mL^−1^ after 72 h (Fig. 1a). *P*. *capeferrum* WCS358 grew similarly in 10% vinasse and LB medium attaining 2.50·10^9^ CFU mL^−1^ after 24 h (Fig. 1b). In contrast, CFU mL^−1^ counts decreased in vinasse 20% during the first 24 h, but then slowly increased reaching 4.03·10^8^ CFU mL^−1^ after 72 h (Fig. 1b).

**Figure 1.**
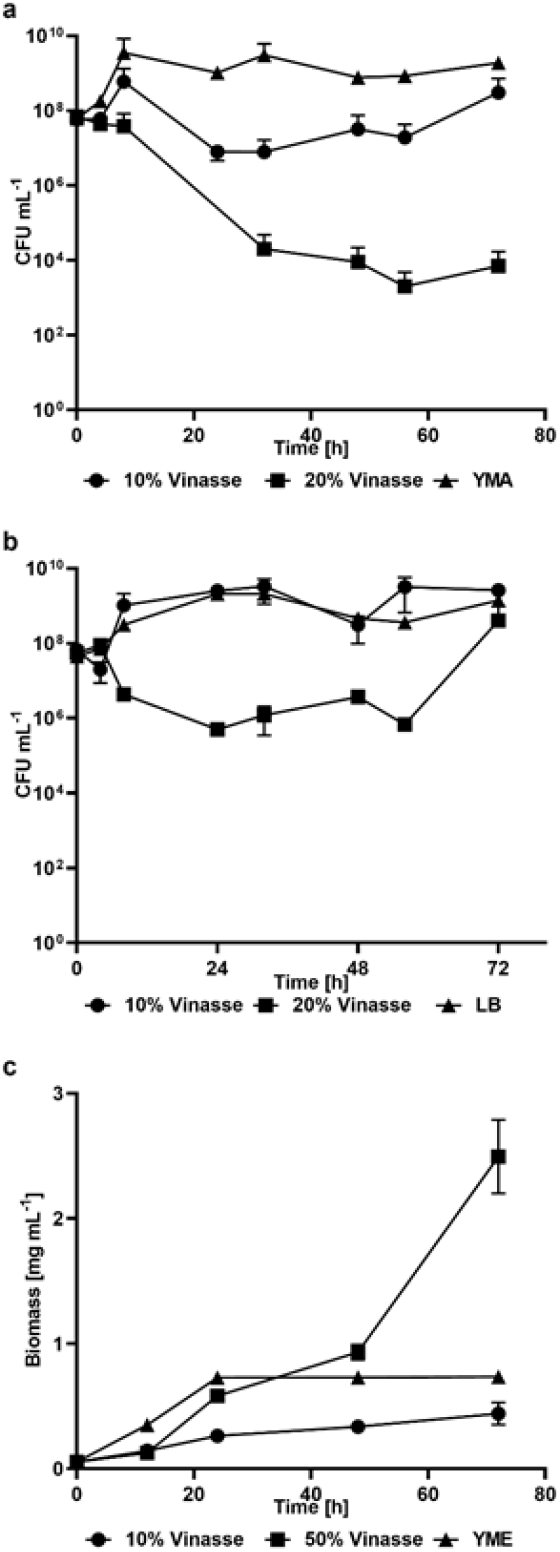
Vinasse as medium for the pure culture of plant-growth promoters. *Rhizobium* sp. N21.2 (a) and *P*. *capeferrum* WCS358 (b) were grown in 10% vinasse and 20% vinasse, and compared to growth in YMA and LB broth, respectively. *T*. *harzianum* MT2 (c) was grown in 10% vinasse and 50% vinasse and compared to growth in YME broth. Error bars represent standard deviations.

*T*. *harzianum* MT2 tolerated higher vinasse concentrations in comparison to both bacteria, even resisting 100% pure vinasse (Suppl. Fig.2c). Two dilutions were then utilized to compare the fungal growth to that in YME broth: 10% vinasse, in which bacteria showed the better growth, and 50% vinasse that allowed the better fungal growth. Growth in 10% vinasse was slower than in YME, in which the stationary phase was attained after 24 h with a biomass of 0.73 mg mL^−1^ (Fig. 1c). In contrast, biomass in 10% vinasse did not exceed 0.44 mg mL^−1^. After a 12 h-lag phase, *T*. *harzianum* MT2 growth markedly increased in 50% vinasse reaching biomass values of 2,49 mg mL^−1^, more than three folds higher than in YME (Fig. 1c).

### 3.2 Vinasse as growth medium for sequential co-cultures

In order to increase the utilization of vinasse as culture medium and to lower the use of clean water for its dilution, co-cultures were evaluated following a two-stage cultivation sequential strategy, as described in Materials and Methods section (Suppl. Fig. 1). Growths were evaluated during the second stage after the inoculation of *T*. *harzianum* MT2 and the supplementation with vinasse. No differences were found between the growth of *T. harzianum* MT2 in single and in co-cultures with *P*. *capeferrum* WCS358, which attained 3.64 mg mL^−1^ and 3.30 mg mL^−1^, respectively, 48 h after vinasse supplementation. Fungal biomasses then decreased to 2.53 mg mL^−1^ and 1.99 mg mL^−1^ (Fig. 2a). Growth of *T. harzianum* MT2 was lower in co-culture with *Rhizobium* sp. N21.2 attaining 1.64 mg mL^−1^ after 48 h of supplementation, but with values of 3.12 mg mL^−1^ after 72 h (Fig. 2a).

**Figure 2.**
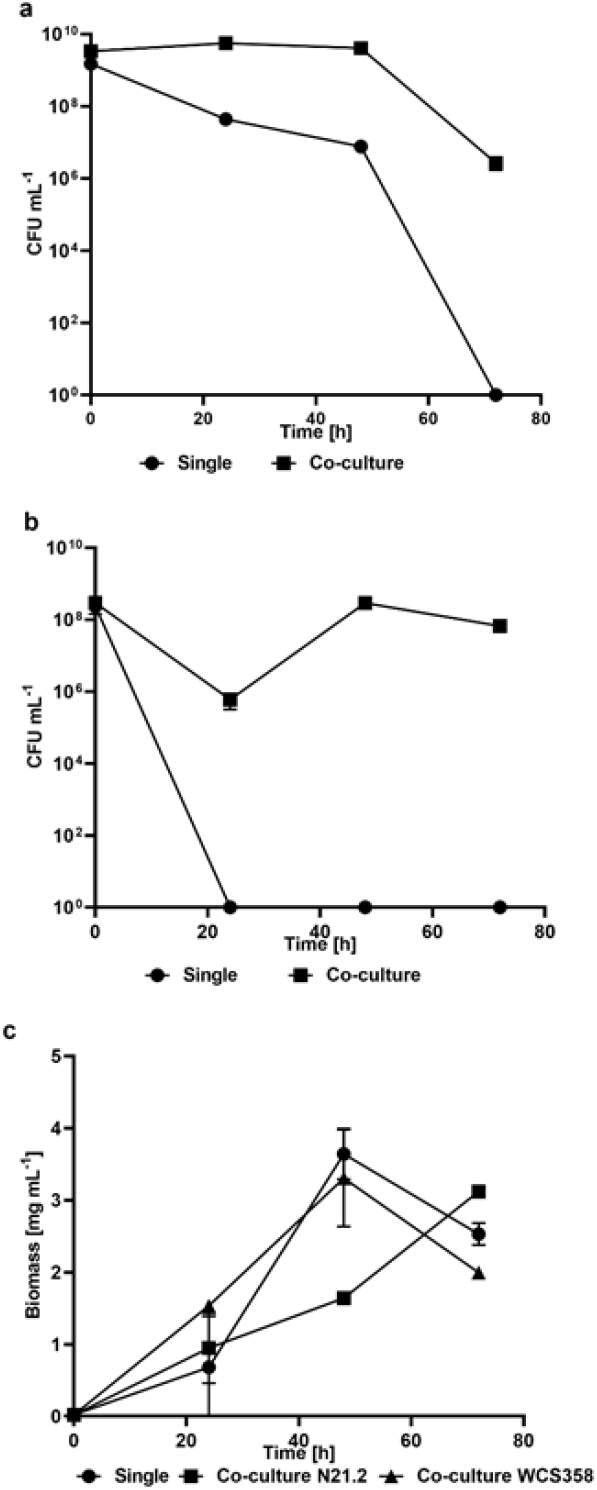
Vinasse as medium for the sequential co-culture of plant-growth promoters. Growth of *Rhizobium* sp. N21.2 (a) and *P*. *capeferrum* WCS358 (b) after the addition of *T. harzianum* MT2 and the supplementation with vinasse, in comparison with single cultures after vinasse supplementation. Growth of *T. harzianum* MT2 (c) after been added to *Rhizobium* sp. N21.2 or *P*. *capeferrum* WCS358 single cultures, in comparison to single culture. Error bars represent standard deviations.

After a decrease of two log units, in presence of *T. harzianum* MT2 and a concentration of 50% vinasse, *P*. *capeferrum* WCS358 rapidly recovered attaining similar values (2.90·10^8^ CFU mL^−1^) to the initials when the culture was supplemented (Fig. 2b). In single cultures only supplemented with vinasse, no colonies of *P*. *capeferrum* WCS358 were obtained after 24 h (Fig. 2b). In contrast, *Rhizobium* sp. N21.2 remained constant at 10^9^ CFU mL^−1^ for 48 h before it decreased to 2.70· 10^6^ CFU mL^−1^ in the next 24 h (Fig. 2c). When *Rhizobium* sp. N21.2 was alone in 50% vinasse, CFU mL^−1^ constantly declined. After 72 h, no colonies were obtained (Fig. 2c).

### 3.3 Microbial growth enhances the characteristics of residual vinasse

The characteristics of the residual vinasses obtained from the single culture of *T. harzianum* MT2 and the sequential co-cultures were analyzed and compared with 50% vinasse (control vinasse). *T. harzianum* MT2 and *P. capeferrum* WCS358+*T. harzianum* MT2 decreased the acidity of vinasse with respect to the control reaching values close to neutrality (pH=6.47 and pH=6.90, respectively). Lower values were obtained with vinasse from co-culture of *Rhizobium* sp. N21.2+*T. harzianum* MT2 (pH=6.08) (Table 1). Vinasse conductivities also diminished in comparison to the control (9.47 dS m^−1^), with significant differences with the single culture of *T. harzianum* MT2 (9.22 dS m^−1^). No differences were observed between both co-cultures (Table 1). Cultivation also reduced the dissolved solids in all residual vinasses in comparison to control (4 °Bx) (Table 1). More marked values were obtained with the single culture of *T. harzianum* MT2 and with the co-culture *P. capeferrum* WCS358+*T. harzianum* MT2 (2,13 °Bx) (Table 1).

**Table 1.** Characterization of vinasses^a^

|  | H<sub>2</sub>O | Control Vinasse | *T. harzianum* MT2 | *P. capeferrum* WCS358<br>+*T. harzianum* MT2 | *Rhizobium* sp. N21.2<br>+*T. harzianum* MT2 |
| --- | --- | --- | --- | --- | --- |
| pH |  | 4.73±0.04 | 6.47±0.14* | 6.90±0.21* | 6.08±0.13* |
| Conductivity [dS m<sup>-1</sup>] |  | 9.49±0.02 | 9.22±0.10* | 9.34±0.08 | 9.36±0.0929 |
| Dissolved solids [°Bx] |  | 4±0 | 2.13±0.11* | 2.13±0.15* | 2.63±0.231* |
| **Vinasse toxicity** |  |  |  |  |  |
| Germination [%] | 100±20* | 19.35±3.22 | 20.43±8.12 | 23.60±6.71 | 17.74±2.28 |
| Hypocotyl [cm] | 2.65±0.36* | 1.40±0.54 | 2.49±0.55* | 2.40±0.75* | 2.66±0.48* |
| Radicle [cm] | 4.00±0.86* | 0.71±0.29 | 1.11±0.20* | 0.92±0.46 | 0.98±0.32 |

<sup>a</sup> Mean values are presented with ±Standard Deviation Values significantly different to the control vinasse (ANOVA followed by Dunnett’s post hoc test at *P*<0.05) are indicated with \*.

Considering that preliminary assays of germination of lettuce seeds with pure vinasses, samples were first diluted 1:5 in water before toxicity tests were performed. No differences were observed in the germination percentages with vinasse from single culture of *T. harzianum* MT2, in comparison to the control vinasse. Slightly higher values were measured with vinasse from *P. capeferrum* WCS358+*T. harzianum* MT2 (23.60%), but lower with *Rhizobium* sp. N21.2+*T. harzianum* MT2 (17.74%). Hypocotyl lengths were longer with *T. harzianum* MT2 (2.49 cm) in comparison to control (1.40 cm). Similar values to the single culture were obtained with *P. capeferrum* WCS358+*T. harzianum* MT2 (2.40 cm), and longer (2.66 cm) with *Rhizobium* sp. N21.2+*T. harzianum* MT2 (Table 1). Length of radicles tended to increase with residual vinasse from *T. harzianum* MT2 (1.11 cm) with respect to the control vinasse (0.71 cm) (Table 1), and with no significant differences between co-cultures.

The overall morphology of the seedlings showed the clearest difference in the toxicity. Seedlings obtained with residual vinasses developed morphologies similar to water treatment. In contrast, seedlings with control vinasse were abnormal with twisted forms (Fig. 3).

**Figure 3.**
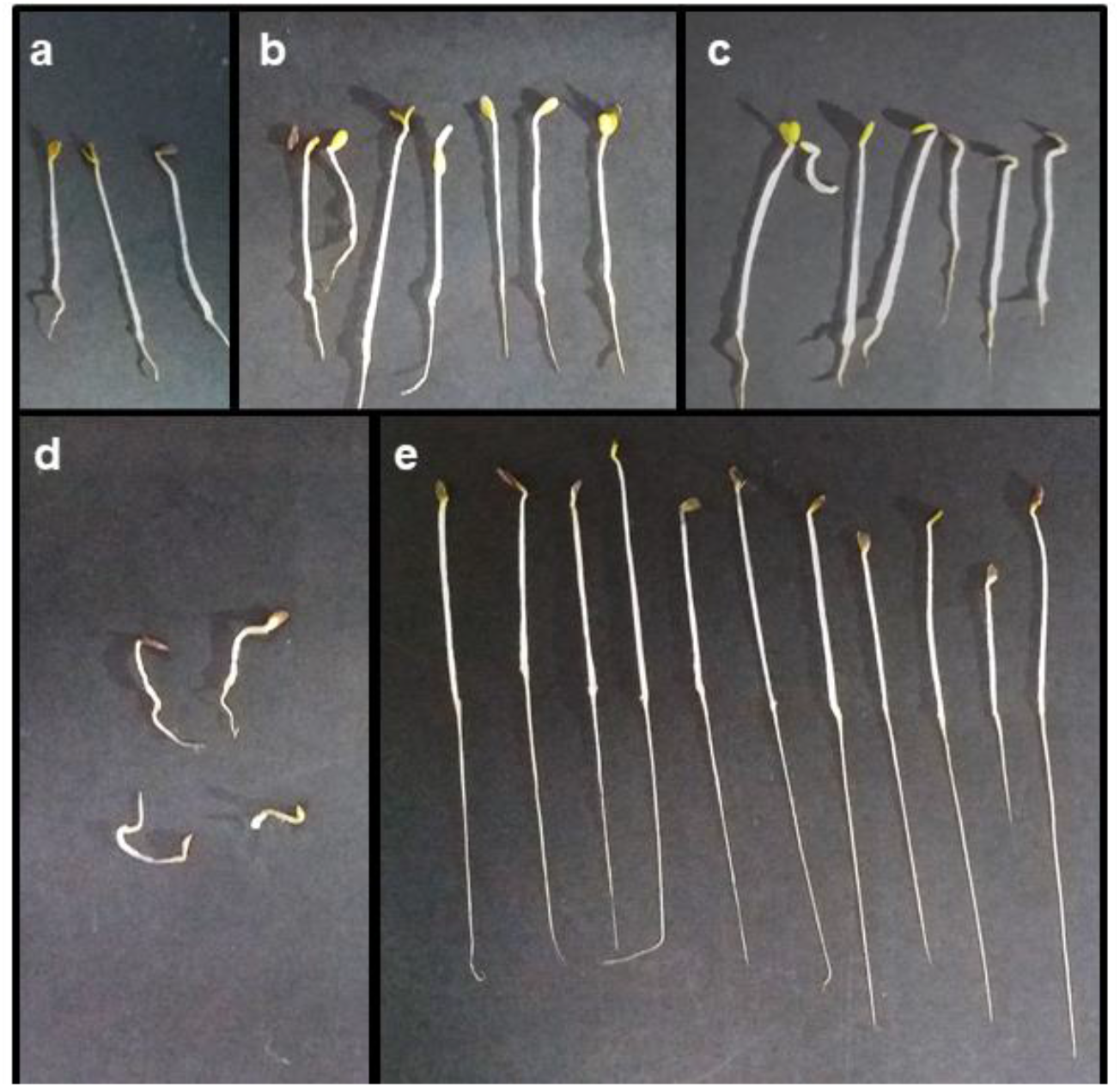
Morphologies of lettuce seedlings. Seeds were germinated in residual vinasses from single culture of *T. harzianum* MT2 (a), sequential co-cultures of *P. capeferrum* WCS358+*T. harzianum* MT2 (b) and *Rhizobium* sp. N21.2+*T. harzianum* MT2 (c) are compared with control vinasse (d) and water (e).

### 3.4 Fertigation with residual vinasses is better than with control vinasse for physical-chemical and toxicity characteristics of soil

Soil samples were fertigated with residual vinasses obtained from single fungal culture and from co-cultures, and then the short-term impact in the physical-chemical and toxicity characteristics were determined. The pH of soils fertigated with residual vinasses were similar (pH 7.13-7.19) to that with control vinasse (pH=7.12), and slightly higher than with water (pH=6.82) (Table 2). The fertigation with vinasses augmented the conductivity in comparison to the irrigation with water (0.35 dS m^−1^). However, in comparison to control vinasse (0.67 dS m^−1^), treatments with residual vinasses from *T. harzianum* MT2 and *Rhizobium* sp. N21.2+*T. harzianum* MT2 showed lower values (0.63 dS m^−1^), and even lower with *P. capeferrum* WCS358+*T. harzianum* MT2 (0.59 dS m^−1^). Other relevant characteristics included an increase in total N, in particular with *P. capeferrum* WCS358+*T. harzianum* MT2, and in the CEC with *T. harzianum* MT2 and *P. capeferrum* WCS358+*T. harzianum* MT2 (Table 2).

**Table 2.** Physical-chemical and chemical characterization and toxicity of soils after treatments with vinasses or water.

|  | H<sub>2</sub>O | Control Vinasse | *T. harzianum* MT2 | *P. capeferrum* WCS358<br>+*T. harzianum* MT2 | *Rhizobium* sp. N21.2<br>+*T. harzianum* MT2 |
| --- | --- | --- | --- | --- | --- |
| pH | 6.82±0.05* | 7.12±0.02 | 7.13±0.04 | 7.19±0.02 | 7.17±0.08 |
| Conductivity [dS m<sup>-1</sup>] | 0.35±0.07* | 0.67±0.08 | 0.63±0.16 | 0.59±0.06 | 0.63±0.13 |
| CO<sub>3</sub><sup>2-</sup> [%] | <0.20 | <0.20 | <0.20 | <0.20 | <0.20 |
| TOC [%] | 3.86 | 3.94 | 4.01 | 4.16 | 4.13 |
| Total N [%] | 0.241 | 0.247 | 0.251 | 0.260 | 0.258 |
| Available P [ppm] | 13.5 | 13.0 | 13.2 | 13.4 | 13.4 |
| CEC [mEq 100 g<sup>-1</sup>] | 16.10 | 15.41 | 16.43 | 16.22 | 16.07 |
| Exchange Cations |  |  |  |  |  |
| Ca<sup>2+</sup> [mEq 100 g<sup>-1</sup>] | 10.50 | 10.56 | 10.55 | 10.74 | 10.71 |
| Mg<sup>2+</sup> [mEq 100 g<sup>-1</sup>] | 2.95 | 3.10 | 4.15 | 3.94 | 3.78 |
| K<sup>+</sup> [mEq 100 g<sup>-1</sup>] | 1.28 | 1.30 | 1.28 | 1.28 | 1.26 |
| Na<sup>+</sup> [mEq 100 g<sup>-1</sup>] | <0.25 | <0.25 | <0.25 | <0.25 | <0.25 |
| **Soil toxicity** |  |  |  |  |  |
| Germination [%] | 98.89±1.96 | 96.63±3.89 | 94.38±3.37 | 97.75±3.37 | 91.01±12.15 |
| Hypocotyl [cm] | 2.18±0.42* | 2.42±0.39 | 2.45±0.44 | 2.45±0.53 | 2.06±0.45* |
| Radicle [cm] | 2.34±0.82 | 2.61±0.69 | 2.63±0.67 | 2.22±0.80* | 2.24±0.67* |

<sup>a</sup> Mean values are presented with ±Standard Deviation
Values of pH, conductivity and soil toxicity significantly different to the control vinasse (ANOVA followed by Dunnett's post hoc test at *P*<0.05) are indicated with \*.

No significant differences were observed in the percentages of germination when soil toxicity was tested in lettuce seeds, with values between 93.3% (fertigation with vinasse from *T. harzianum* MT2 culture) and 98% (irrigation with water) (Table 2). Interestingly, hypocotyl lengths were higher with control vinasse (2.42 cm) and with vinasses from *T. harzianum* MT2 and *P. capeferrum* WCS358+*T. harzianum* MT2 cultures (2.45 cm), that the irrigation with water (2.18 cm). Radicle lengths were higher with control (2.61 cm) and with vinasse from *T. harzianum* MT2 culture (2.63 cm). In contrast, the shortest lengths were obtained with *Rhizobium* sp. N21.2+*T. harzianum* MT2 and *P. capeferrum* WCS358+*T. harzianum* MT2: 2.23 cm and 2.22 cm, respectively (Table 2).

### 3.5 Fertigation with residual vinasses improves biological characteristics of soil

Quantification of enzymatic activities in soils fertigated with residual vinasses showed that UA after treatment with vinasses from *T. harzianum* MT2 culture (21.08 μg N-NH_4_ g^−1^ h^−1^) was lower than with control vinasse (27.62 μg N-NH_4_ g^−1^ h^−1^). Even lower values were determined with *P*. *capeferrum* WCS358+*T. harzianum* MT2 (19.73 μg N-NH_4_ g^−1^ h^−1^), very similar to water (19.30 μg N-NH_4_ g^−1^ h^−1^). Intermediate activity (24.56 μg N-NH_4_ g^−1^ h^−1^) was determined with *Rhizobium* sp. N21.2+*T. harzianum* MT2 (Fig. 4a). Residual and control vinasses induced similar AP activities in comparison to water. The exemption was the vinasse from *P*. *capeferrum* WCS358+*T. harzianum* MT2 co-cultures (213.56 μg g^−1^ h^−1^), which caused a reduction in comparison to water (291.63 μg g^−1^ h^−1^) and to control (268.69 μg g^−1^ h^−1^) (Fig. 4b). No differences were determined in FDA and CA, regardless of the irrigation used (Fig. 4c and d). Heterotrophic microbial population after fertigation with residual vinasses was also not modified significantly, except for vinasse from *Rhizobium* sp. N21.2+*T. harzianum* MT2 co-culture, attaining 1,96·10^7^ CFU g^−1^, higher than with control vinasse (9,32·10^6^ CFU g^−1^) and water (8,15·10^6^ CFU g^−1^) (Fig. 4e).

**Figure 4.**
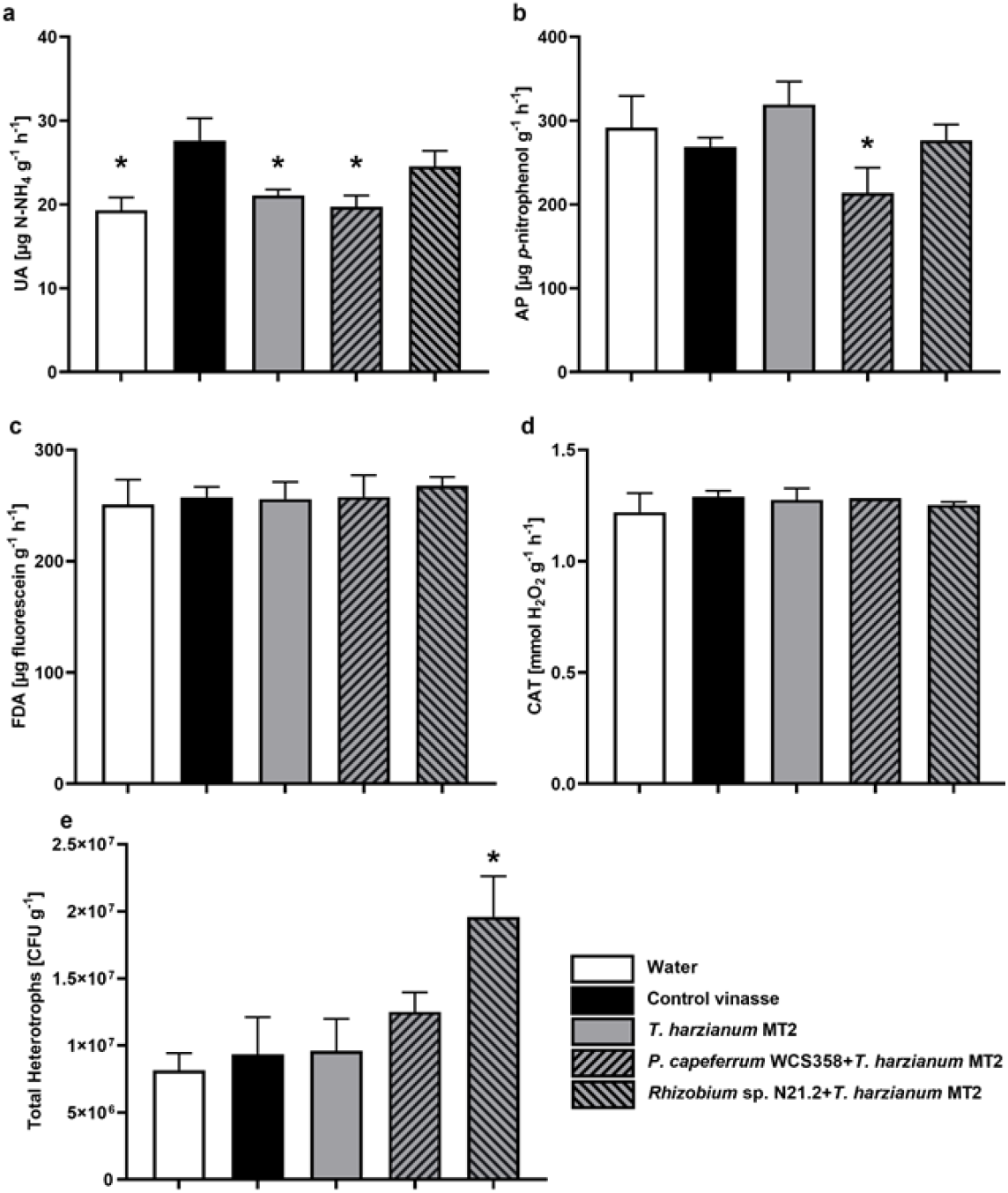
Biological characteristics of fertigated soil. Urease activities (a), acid phosphatase (b), fluorescein diacetate hydrolysis (c), catalase (d) activities, and heterotrophic microorganisms (e) were determined after irrigation with water or fertigation with control vinasse, residual vinasses from single culture of *T*. *harzianum* MT2, sequential co-cultures of *P. capeferrum* WCS358+*T. harzianum* MT2 and *Rhizobium* sp. N21.2+*T. harzianum* MT2. Error bars represent standard deviations. Values significantly different from the control vinasse (ANOVA followed by Dunnett’s post hoc test, *P*<0.05) are indicated with *.

The metabolic diversity of the microbial community in the fertigated soils was assessed using Biolog Ecoplates. The evaluation of the Average Metabolic Response (AMR) showed two groups of treatments that differed from water irrigation. One group, including the treatment with vinasses from *T. harzianum* MT2 culture and from *P*. *capeferrum* WCS358+*T. harzianum* MT2 co-culture, showed slower increases in the AMR with maximal values of 0.20 and 0.18 after 120 h, respectively (Fig. 5a). A second group with a faster increase in the AMR and higher final values (0.22 and 0.23) included the fertigation with control vinasse and vinasses from *T. harzianum MT2+Rhizobium* sp. N21.2 co-cultures. Soil irrigated with water showed the slowest increase in AMR throughout this study, presenting final values of 0.13 (Fig. 5a).

**Figure 5.**
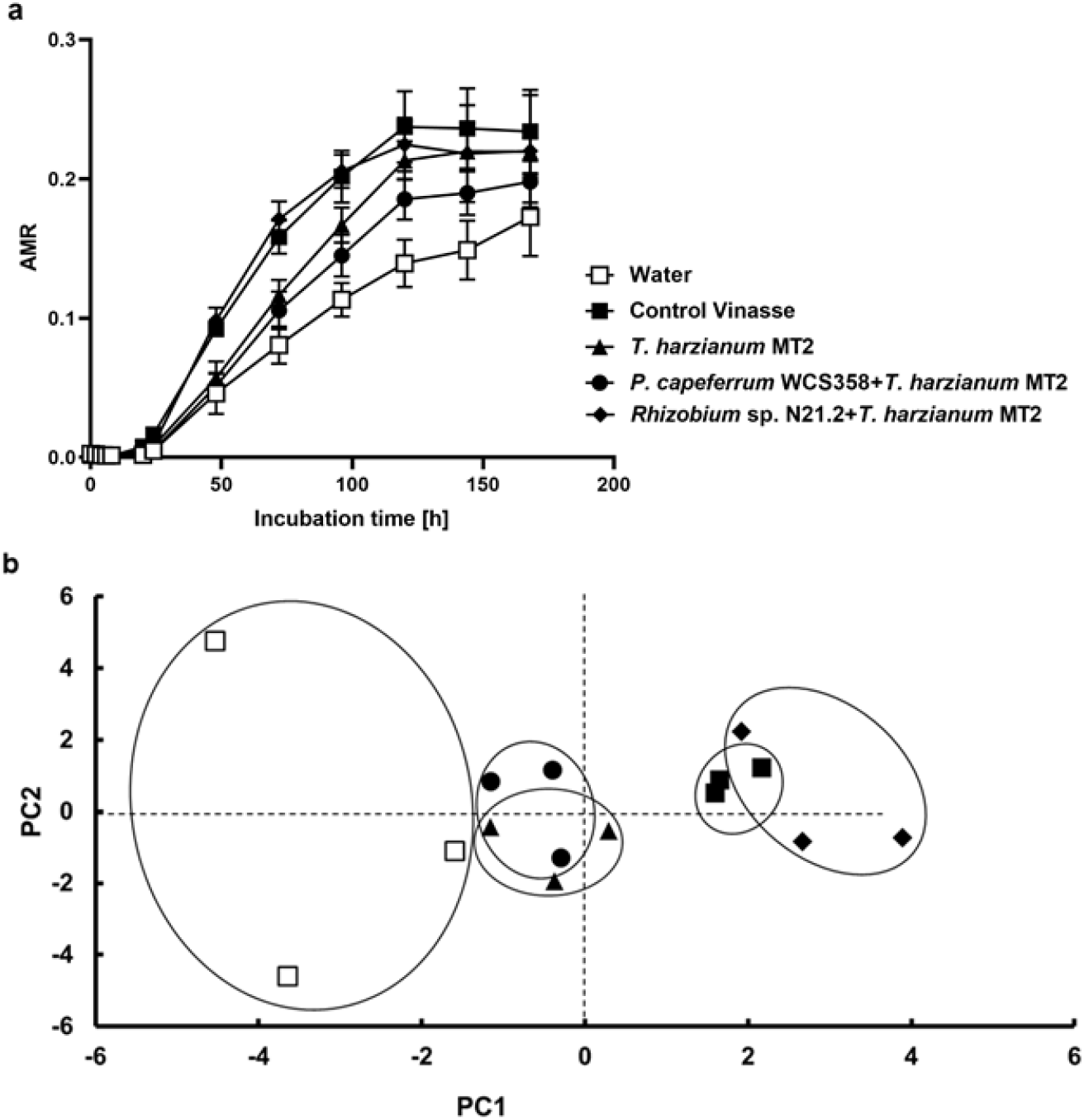
Metabolic diversity of fertigated soils. Biolog EcoPlates were utilized to evaluate the Average Metabolic Response (AMR) of soils after irrigation with water or fertigation with control vinasse or residual vinasses. Values of AMR were obtained at different incubation times of the Ecoplates (a). Principal Component Analysis (b) was performed with data obtained after 72 h of incubation.

Employing the Principal Component Analysis, the separation on the PC1 axis in the two groups mentioned above could be clearly distinguished, though no major differences were observed on the PC2 axis (Fig. 5b). The first group (residual vinasses from *T. harzianum* MT2 and *P*. *capeferrum* WCS358+*T. harzianum* MT2) together with water irrigation was located towards the negative PC1 values, due to the influence of the carbon sources Glucose-1-phosphate, D-Xylose and L-Phenylalanine (Fig. 5b). Within this group, the most distant and dispersed treatment was the soil irrigated with water. The second group (control vinasse and vinasse from *Rhizobium* sp. N21.2+*T. harzianum* MT2 co-culture), on the contrary, was located towards the positive side of PC1, mainly due to the utilization of D-Mannitol, L-Arginine and L-Asparagine (Fig. 5b).

## 4 Discussion

The main limitation for the exploitation of vinasse as culture medium is the toxicity of this by-product. Results presented in this report show that *T. harzianum* can be cultured in sugarcane vinasse for fungal mass production with better results than a common culture medium. The tolerance of *T. harzianum* MT2 is in agreement with the reported high fungal tolerance to vinasse (Rodrigues Reis et al., 2020). Inhibition of microbial growth in vinasse, mainly attributed to toxic phenolics like luteolin, tricin, apigenin, and naringenin (Rodrigues Reis et al., 2020), could explain the lower growth of *T. harzianum* MT2 in 100%. The analysis of the fungal growth suggests that *T. harzianum* MT2 first utilizes readily metabolizable nutrients, and then compounds derived from complex components. In agreement, when *Aspergillus niger* grows in vinasse, monosaccharides are first accumulated from the degradation of complex molecules with a relatively weak increase in biomass. After glucose and fructose exhaustion, *A. niger* utilizes mannitol with a sudden increase in the biomass (Chuppa-Tostain et al., 2018). From a perspective of circular economy, these results indicate the possibility of exploiting sugarcane vinasse for obtaining the valuable biomass of *T. harzianum* for agricultural practices in an inexpensive manner reducing the production costs related to the culture medium.

Considering the potential of mixed bioinoculants and the possible improvements in the characteristics of the residual vinasse, *P*. *capeferrrum* WCS358 and *Rhizobium* sp. N21.2 were evaluated in sequential co-cultures with *T. harzianum* MT2. Both bacteria showed lower tolerance, but comparable to others (Ventorino et al., 2019). A lack of biphasic growth in *P*. *capeferrrum* WCS358 or *Rhizobium* sp. N21.2 could be related to a minor capacity to degrade complex compounds, which contrasts with the recognized lytic enzyme production in *T. harzianum* (Schuster and Schmoll, 2010). Despite this low tolerance, *P*. *capeferrrum* WCS358 and *Rhizobium* sp. N21.2 growths in 10% vinasse were similar than in the corresponding laboratory culture media and survived to higher concentrations in co-cultures. This enhancement could be related to the degradation of phenolics by *T. harzianum* MT2. In agreement, previous reports show that co-cultures of bacteria and fungi have advantages in terms of resistance to toxic compounds (Losa and Bindschedler, 2018). These results are relevant since they indicate that the biomass for the first stage of sequential co-cultures can also be produced in vinasse allowing a higher utilization of vinasse and lower costs of production.

From a perspective of a green economy, the characteristics of the wastes generated after utilizing vinasse as culture medium, and their ecological impact should also be considered. Several studies already reported that microbial growth in vinasse decreases its toxicity (Ahmed et al., 2022; Rulli et al., 2020). The neutralization, for instance, is a valuable feature, considering that that the acidity is a serious concern for fertigation. The lower acidity with residual vinasse from *T. harzianum* MT2 could be related to the consumption of organic acids generated from the metabolism of carbohydrates, as reported previously for *A. niger* (Chuppa-Tostain et al., 2018). The even higher pH in vinasse from *P*. *capeferrum* WCS358+*T. harzianum* MT2 co-culture could also be related to the well-known utilization of organic acids by *Pseudomonas* spp. (Lynch and Franklin, 1978). Both cases are in agreement with the decreases in the amount of dissolved solids determined in the corresponding residual vinasses. Rhizobia also utilize organic acids with a preference over carbohydrates (Iyer et al., 2016). However, the survival of *Rhizobium* sp. N21.2 was reduced in 50% vinasse, even in presence of *T. harzianum* MT2, which could explain the lower neutralization and the higher amount of dissolved solids in these co-culture. The probable degradation of phenolics by *T. harzianum* MT2 in both single and co-culture could also explain the decrease in vinasse toxicity evidenced with lettuce seeds. Other factors, like the decrease in the conductivity of residual vinasses, cannot be ruled out. For instance, high concentrations of soluble salts have also been implicated in the phytotoxicity of effluents (Wang, 1991).

The analysis of the soils also shows that the environmental impact of the residual vinasse from *T. harzianum* MT2 is lower than with control vinasse. The analysis of soil conductivity after fertigation with residual vinasses, in comparison to the irrigation with water, indicate a salinization process, possibly due to the higher Mg^2+^ content. However, values of conductivities were similar to that of control vinasse and, though not statistically significant, lower after fertigation with residual vinasse from *P*. *capeferrum* WCS358+*T. harzianum* MT2. Soil toxicity parameters diminished, particularly evidenced in the hypocotyl and radicle lengths, with co-cultures. It is plausible that *P*. *capeferrum* WCS358 and *Rhizobium* sp. N21.2 produce plant-growth regulators or protect the germinated seeds from the stress generated by the vinasse.

Other biological parameters also showed that fertigation with vinasse from *T. harzianum* MT2 culture is less detrimental to soil quality than fertigation with raw control vinasse. Urease activity (UA) is a relevant indicator of soil quality, mainly related to the N cycle and highly influenced by soil disturbances (Adetunji et al., 2020). For instance, sewage sludge increase UA causing the release of nitrogen available for plants. The comparison of UA and total N values (Table 2) suggest that residual vinasse from *T. harzianum* MT2 contributes to the overall N cycle lowering the enzymatic activity and supplying N to the soil. To note, the efficiency is better with vinasse from *P*. *capeferrum* WCS358+*T. harzianum* MT2 co-culture. Phosphatases participate in the P cycle releasing phosphate from phosphate monoester that can be biologically uptaken (Adetunji et al., 2020). Vinasse from *T. harzianum* MT2 has a positive impact on the AP activity. However, the decrease after fertigation with vinasse from co-cultures, in particular from *P*. *capeferrum* WCS358+*T. harzianum* MT2, suggests a negative effect on P cycle. In sights of a putative fertigation with residual vinasses from co-cultures, these results suggest that proper amendments would be required, even if the values of available P in soil were higher that with control vinasse. The constant values of the fluorescein diacetate hydrolysis, together with the increase in the heterotrophic microorganisms after fertigation with vinasse from *Rhizobium* sp. N21.2+*T. harzianum* MT2 indicate that this treatment decrease, in relative terms, the microbial activity in soil (Green et al., 2006).

The physiological profiling of the microbial communities after fertigation with *T. harzianum* MT2 also showed an improvement, which was enhanced with vinasse from *P*. *capeferrum* WCS358+*T. harzianum* MT2 co-culture. In contrast, AMR with vinasse from *Rhizobium* sp. N21.2+*T. harzianum* MT2 was closer to the control vinasse treatment. *P*. *capeferrum* WCS358 and *T. harzianum* MT2 may produce beneficial compounds with a positive effect on the soil. However, it should also be taken into account the presence of fungal propagules and bacterial cells in the residual vinasses. To note, in an approach to industrial developments that require non-expensive methods, in this work residual vinasses were not sterilized before fertigation.

## 5 Conclusions

The actual vision considers vinasse as a by-product with broad potential. Vinasse can be utilized for the production of biomass valuable for agricultural practices. Toxicity can be diminished by the fungal growth, allowing safer fertigation. The sequential co-culture with plant-growth promoting bacteria permits to obtain a mixed bioinoculant enhancing the characteristics of the residual vinasse. However, it is important to properly select the bacterium for the co-culture. These results are relevant in terms of circular and green economy considering that an agroindustrial by-product can be utilized for the production of inoculants for agriculture, generating residual vinasse of lower ecological impact.

## Acknowledgement

This work was supported by the Consejo Nacional de Investigaciones Científicas y Técnicas (CONICET, PIP 2015-0946, PIP 2021-2436, PUE 22920160100012CO), Agencia Nacional de Promoción Científica y Tecnológica (PICT 2016 N° 0532; PICT 2019 N° 03336, PICT 2016 N° 2013, PICT 2018 N°3500, PICT 2018 N°01765), and Secretaría de Ciencia, Arte e Innovación Tecnológica from the Universidad Nacional de Tucumán (PIUNT D609).

---

**Supplementary Table 1.** Characteristics of sugarcane vinasse utilized in this work

| Determination | Value |
| --- | --- |
| pH | 4,9 |
| Volatile solids | 41350 mg L<sup>-1</sup> |
| Total solids | 62480 mg L<sup>-1</sup> |
| Fixed solids | 21130 mg L<sup>-1</sup> |
| Dissolved solids | 7,20 °Bx |
| Organic matter | 39283 mg L<sup>-1</sup> |
| Conductivity | 21,7 mS cm<sup>-1</sup> |
| Total P | 250 mg P L<sup>-1</sup> |
| Total N | 0,08 % |
| Cl<sup>-</sup> | 1974 mg Cl L<sup>-1</sup> |
| S<sup>2-</sup> | 32 mg S L<sup>-1</sup> |
| Ca<sup>2+</sup> | 1,0 g Ca L<sup>-1</sup> |
| Mg<sup>2+</sup> | 376 mg L<sup>-1</sup> |
| Na<sup>+</sup> | 128 mg L<sup>-1</sup> |
| K<sup>+</sup> | 10 g L<sup>-1</sup> |

**Supplementary Figure 1.**
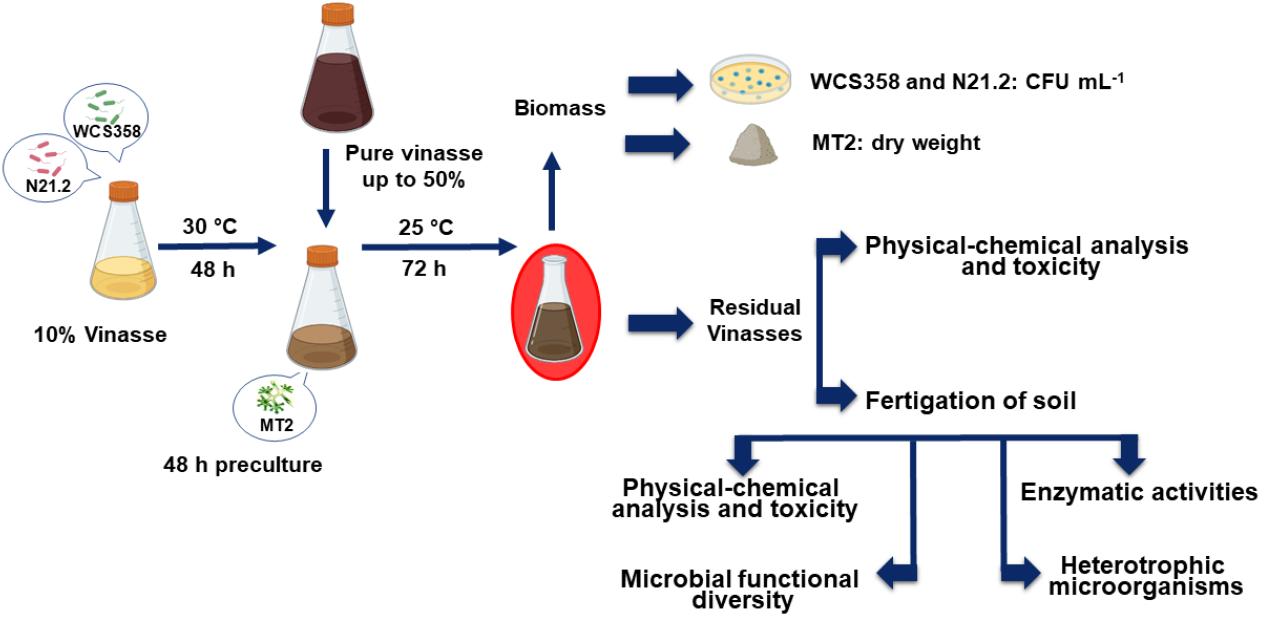
Sequential co-cultures of *T. harzianum* with bacteria in vinasse. *Rhizobium* sp. N21.2 and *P*. *capeferrum* WCS358 were grown 48 h in 10% vinasse at 30 °C. *T. harzianum* MT2 was then inoculated together with pure vinasse up to a final concentration of 50%. Cultures were continued at 25 °C for 72. During these 72 h, biomass was determined with CFU mL-1 for bacteria or dry weight for *T. harzianum* MT2. Physical-chemical analysis and toxicity were determined in residual vinasses. Fertigation of soils was also performed with residual vinasses.

**Supplementary Figure 2.**
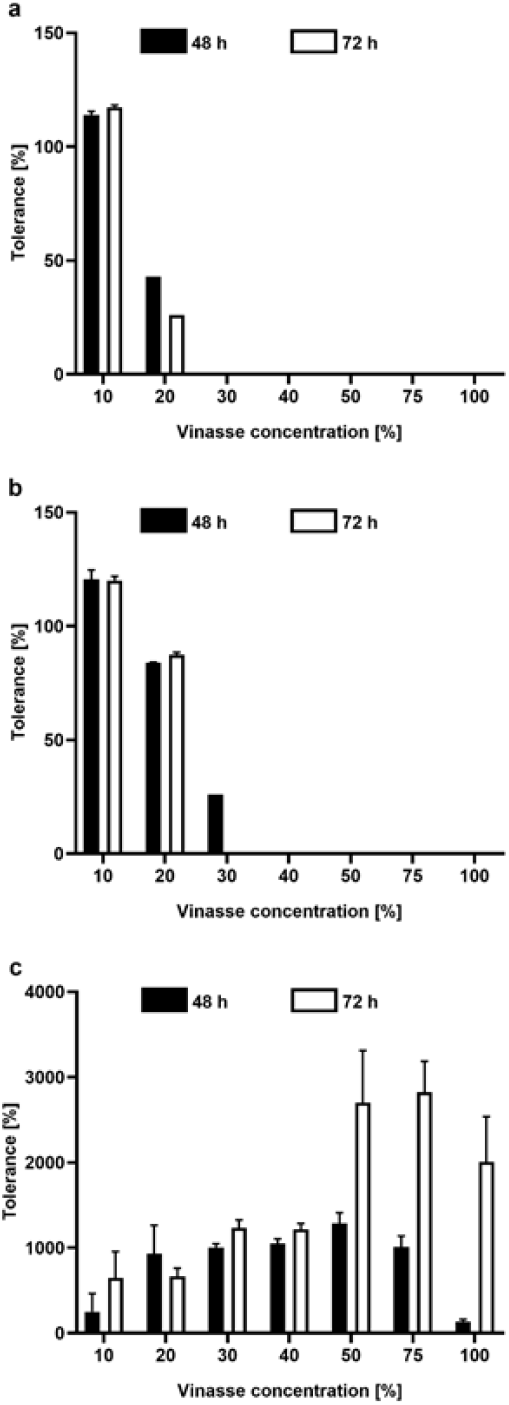
Tolerance to increassing concentrations of vinasse. After inoculations, *Rhizobium* sp. N21.2 (a), *P*. *capeferrum* WCS358 (b) and *T*. *harzianum* MT2 (c) were incubated for 48 h (black bars) and 72 h (white bars). Bacterial and fungal growth was determined through the quantification of CFU mL^−1^ and mg mL^−1^, respectively, and expressed as % of the corresponding inoculum. Error bars represent standard deviations.

